# Predicting the pathway involvement of metabolites annotated in the MetaCyc knowledgebase

**DOI:** 10.1101/2024.10.29.620954

**Authors:** Erik D. Huckvale, Hunter N.B. Moseley

## Abstract

The associations of metabolites with biochemical pathways are highly useful information for interpreting molecular datasets generated in biological and biomedical research. However, such pathway annotations are sparse in most molecular datasets, limiting their utility for pathway level interpretation. To address these shortcomings, several past publications have presented machine learning models for predicting the pathway association of small biomolecule (metabolite and zenobiotic) using data from the Kyoto Encyclopedia of Genes and Genomes (KEGG). But other similar knowledgebases exist, for example MetaCyc, which has more compound entries and pathway definitions than KEGG. As a logical next step, we trained and evaluated multilayer perceptron models on compound entries and pathway annotations obtained from MetaCyc. From the models trained on this dataset, we observed a mean Matthews correlation coefficient (MCC) of 0.845 with 0.0101 standard deviation, compared to a mean MCC of 0.847 with 0.0098 standard deviation for the KEGG dataset. These performance results are pragmatically the same, demonstrating that MetaCyc pathways can be effectively predicted at the current state-of-the-art performance level.

**Author summary:** Many thousands of different molecules play important roles in the processes of life. To generally handle the complexity of life, biological and biomedical researchers typically organize the molecular parts and pieces of biological processes into pathways of biomolecules and their myriad of molecular interactions. While the role of large macromolecules like proteins are well characterized within these pathways, the role of small biomolecules are not as comprehensively known. To close this knowledge gap, several machine learning models have been trained on data from a knowledgebase known as the Kyoto Encyclopedia of Genes and Genomes (KEGG) to predict which pathways a small biomolecule is associated with. More data generally improves these machine learning models. So in this work, we used the MetaCyc knowledgebase to increase the amount of data available by about ten-fold and then trained new machine learning models that demonstrate comparable prediction performance to models trained on KEGG, but covering 8-fold more pathways defined in MetaCyc vs KEGG.

## Introduction

Metabolism is the set of biochemical reactions and related processes for sustaining life. These biochemical reactions convert environmentally-derived and endogenous molecular and energy resources into useful forms of chemical energy and molecular building blocks that drive biological processes, while converting metabolic wastes into forms that can be disposed of. Metabolism can also be viewed as the entirety of these biochemical reactions organized as a metabolic network. Related biological processes can be included into a more general molecular interaction network that is centered on metabolism. Such networks are representable as a hypergraph with biomolecules as nodes and chemical reactions and non-covalent molecular interactions as edges. Certain connected parts or subgraphs of the metabolic-centric molecular interaction network are then defined as distinct “pathways” (1–3).

Compounds connected within a pathway are defined as being associated with that pathway, which is typically represented as a pathway annotation for the compound. Pathway annotations are highly useful for mechanistic interpretation of complex molecular datasets. These annotations are often used in pathway annotation enrichment analysis to identify perturbed pathways based on large datasets of perturbed biomolecules, providing pathway-level insight and interpretation of such datasets. While knowledgebases such as the Kyoto Encyclopedia of Genes and Genomes (KEGG) (4) and MetaCyc (5) contain useful pathway annotations for several thousand compounds, the majority of detected compounds in metabolomics datasets have no pathway annotations, limiting their use in pathway annotation enrichment analysis. Considering the costly and cumbersome work involved in experimentally determining these annotations in vitro, many researchers could benefit from predicting pathway annotations for compounds in silico. Several prior publications present machine learning models that perform this pathway prediction task, trained and tested with data from KEGG (6–9)(10).

Based on the success of these models trained on KEGG-derived datasets, a logical next step is to attempt constructing a similar machine learning dataset from MetaCyc compounds and pathway annotations. The MetaCyc dataset focuses on metabolic pathways as compared to KEGG which contains a wider variety of types of pathways: only 184 out of 502 pathways in KEGG are metabolic pathways. Moreover, MetaCyc contains far more pathway and compound entries. In this work, we demonstrate that MetaCyc pathways can be predicted as effectively as KEGG pathways.

## Materials and methods

The construction of a dataset for model training and evaluation requires the molecular structures of the compounds along with their pathway annotations. Our molecular structure parsing methods expect molfiles (11). On July 9^th^ 2024, we downloaded the molfiles from MetaCyc’s website here https://metacyc.org/download.shtml. The MetaCyc pathways, similar to KEGG, are organized into a hierarchy as seen here: https://metacyc.org/META/class-tree?object=Pathways. The compound to pathway mappings were downloaded on July 9^th^ 2024. We accessed them by traversing the pathway hierarchy using MetaCyc’s web API at the following base URL: https://biocyc.org/META/ajax-direct-subs. With the pathway ID query parameter, you can get the children of a given pathway node. Starting with the base ID ‘Pathways’, the first requestion URL is https://biocyc.org/META/ajax-direct-subs?object=Pathways. Next, you obtain children pathway IDs and recursively request children pathways. Traversing the hierarchy in this way enables access to all the MetaCyc pathway IDs from the root node down to the leaf nodes. Compound associations were obtained from MetaCyc’s web API using this base URL: https://websvc.biocyc.org/apixml?fn=compounds-of-pathway. Again, a pathway ID query parameter is specified. For example, the URL https://websvc.biocyc.org/apixml?fn=compounds-of-pathway&id=META:PWY-7723&detail=low obtains the compounds associated with the pathway corresponding to pathway ID META:PWY-7723.

After obtaining the MetaCyc molfiles and compound-pathway mappings, we constructed the MetaCyc dataset with the methods used for the KEGG dataset (10). This involved creating feature vectors representing compounds, feature vectors representing pathways, and concatenating them together in a cross join. The resulting compound-pathway paired feature vector is then concatenated with a boolean label indicating whether the given compound is associated with the given pathway (8). We created the compound features using an atom coloring technique introduced in our lab (7,12–14). The atom colors represent molecular substructures 1, 2, or 3 bonds away from a central atom node and the feature values are the number of times that substructure appears in the compound. With these atom color counts acting as features for an individual compound, the pathway features are an aggregation of the features of the compounds associated with a given pathway (8). Both the compound features and pathway features were deduplicated and then normalized (10). Table 1 compares the MetaCyc dataset to the KEGG dataset. We see that with nearly 40 million entries, the MetaCyc-derived dataset is the largest published to date, having more than 10-fold the number of paired compound-pathway entries than the KEGG-derived dataset. This is mostly due to the 8-fold larger number of pathway definitions in MetaCyc vs KEGG.

**Table 1.** Comparing the KEGG dataset to the MetaCyc dataset.

| **Dataset** | **#Compounds** | **#Pathways** | **#Compound Features** | **#Pathway Features** | **#Entries** | **Reference** |
| --- | --- | --- | --- | --- | --- | --- |
| **KEGG** | 6,485 | 502 | 16,509 | 11,321 | 3,255,470 | (10) |
| **MetaCyc** | 9,847 | 4,055 | 19,081 | 15,349 | **39,929,585** | Current Study |

With the pathways being hierarchically organized, there are pathways at different hierarchical levels. We will refer to the top-level pathways, e.g. ‘Biosynthesis’, ‘Detoxification’, ‘Glycan Pathways’, ‘Transport’, etc. as level 1 or L1. Likewise, we refer to the level 2 pathways as L2, and the remaining as L3, L4, L5, L6, L7, and L8. To determine the impact of excluding certain pathway levels from the training set on the performance of the remaining pathways, we constructed two subsets of the full MetaCyc dataset i.e. L2+ and L3+. The L1+ dataset contains all of the pathways in the hierarchy, while the L2+ excludes the L1 pathways and contains the L2 pathways and all pathways underneath them in the hierarchy. Likewise, the L3+ dataset excludes the L1 and L2 pathways. Table 2 shows stats for each of these datasets.

**Table 2.** Comparing the subsets of the MetaCyc dataset where L1+ is the full MetaCyc dataset, L2+ excludes L1 pathways, and L3+ excludes L1 and L2 pathways.

| **Dataset** | **#Entries** | **#Pathways** |
| --- | --- | --- |
| **L1+** | 39,929,585 | 4,055 |
| **L2+** | 39,811,421 | 4,043 |
| **L3+** | 38,541,158 | 3,914 |

For our analyses related to the hierarchy levels, we rolled L8 into L7 to form the L7+L8 hierarchy level for more statistically meaningful results, since L8 only had 11 pathways. The reason we did not roll the L1 pathways into L2, is because the L1 pathways are much larger than the L8 pathways, even though there are only 12 L1 pathways. Fig 1 shows the number of pathways in each hierarchy level from L1 to L7+L8. Fig S1 shows this same info, but it shows the number of L7 and L8 pathways separately.

**Fig 1.**
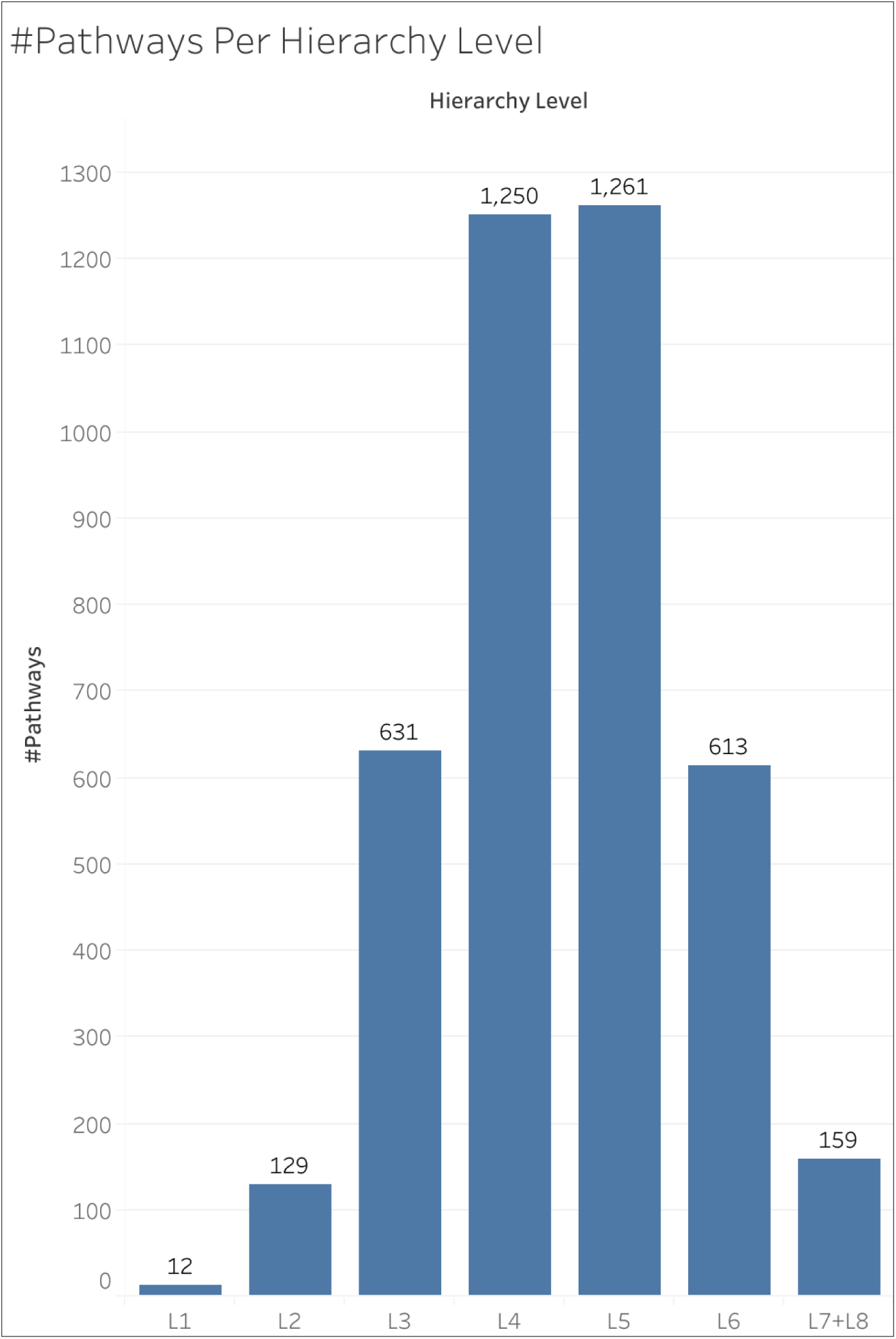
The number of pathways within each hierarchy level.

We used a cross validation (CV) analysis with characteristics of both a bootstrap and jackknife analysis, involving 200 or 50 separate iterations of 10-fold cross validation. On each CV iteration, we split the dataset into a train and test set in a stratified manner across 10 folds (15), ensuring that the proportion of positive entries in the train set is as close as possible to the proportion in the test set. We trained a multi-layer perceptron binary classifier on the train set composed of 9 folds and evaluated on the test set composed of a single fold, collecting the number of true positives (TP), true negatives (TN), false positives (FP), and false negatives (FN). This enabled us to calculate metrics including the Matthews correlation coefficient (MCC) (16) on each CV iteration and then to calculate a mean, median, and standard deviation of each metric across CV iterations. It also enabled us to calculate the MCC per compound and per pathway by counting the TP, TN, FP, and FN in the test set for a given compound or pathway (since each entry is a compound-pathway pair) and then summing those counts across all CV iterations. An MCC cannot be calculated on a single CV iteration since there may not be enough positive entries in the test set for a single compound or pathway to avoid a division by zero. However, we can calculate an overall MCC for an individual compound or pathway by summing TP, TN, FP, and FN across all CV iterations. Likewise, we calculated the MCC for each hierarchy level by summing the TP, TN, FP, and FN across all pathways in a given hierarchy level across all CV iterations. We calculated the overall MCCs of individual compounds and pathways from the L1+ dataset (i.e. the full dataset) and ran the L1+ dataset for 200 CV iterations. We ran the L2+ and L3+ datasets for 50 CV iterations and used the results from those datasets to compare overall MCC across different hierarchy levels. We ran the L1+ dataset for more iterations to ensure a valid MCC calculation for individual compounds or pathways, some of which might not have many positive entries, necessitating more CV iterations to have enough TP or FP.

The hardware used for this work included machines with up to 2 terabytes (TB) of random-access memory (RAM) and central processing units (CPUs) of 3.8 gigahertz (GHz) of processing speed. The name of the CPU chip was ‘Intel(R) Xeon(R) Platinum 8480CL’. The graphic processing units (GPUs) used had 81.56 gigabytes (GB) of GPU RAM, with the name of the GPU card being ‘NVIDIA H100 80GB HBM3’.

All code for this work was written in major version 3 of the Python programming language (17). Data processing and storage was done using the Pandas (18), NumPy (19), and H5Py (20) packages. Models were constructed and trained using the PyTorch Lightning (21) package built upon the PyTorch (22) package. The stratified train test splits were computed using the Sci-Kit Learn package (23). Results were initially stored in an SQL database (24) using the DuckDB (25) package. Results were processed and visualized using Jupyter notebooks (26), the seaborn package (27) built upon the MatPlotLib (28) package, and the Tableau business intelligence application (29).

## Results

### Main results

Table 3 includes the mean, median, and standard deviation of the MCC calculated across CV iterations of the L1+ (full), L2+, and L3+ datasets. The mean and median MCCs are very close, even though distributions of MCC do not look unimodal nor very symmetric. Table S1 shows these results for other metrics i.e. accuracy, precision, recall, F1 score, and specificity. We observe a decrease in performance when we exclude higher level pathways.

**Table 3.** Matthew’s correlation coefficient statistics for models trained on the L1+, L2+, and L3+ datasets.

| Hierarchy Levels Included | Mean MCC | Median MCC | Standard Deviation |
| --- | --- | --- | --- |
| **L1+ Dataset** | 0.8446 | 0.8454 | 0.0101 |
| **L2+ Dataset** | 0.8337 | 0.8347 | 0.0113 |
| **L3+ Dataset** | 0.8199 | 0.8214 | 0.0080 |

Fig 2 shows the distribution of MCC across the 200 CV iterations of the L1+ dataset.

**Fig 2.**
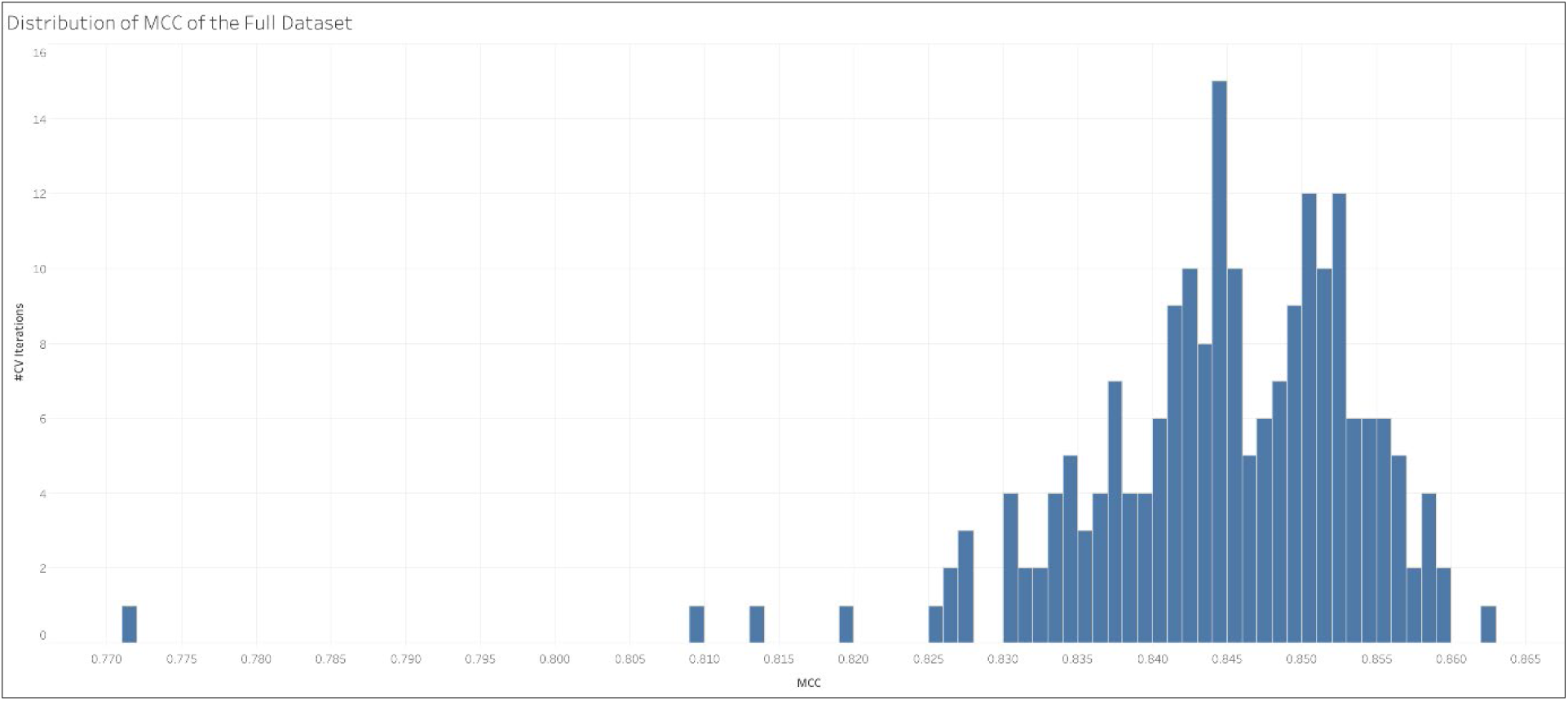
Distribution of the MCC across all CV iterations on the L1+ (full) MetaCyc dataset.

Fig 3 shows the overall MCC across pathways at certain hierarchy levels and across all CV iterations of each dataset i.e. L1+, L2+, and L3+. We see that performance consistently declines at deeper hierarchy levels.

**Fig 3.**
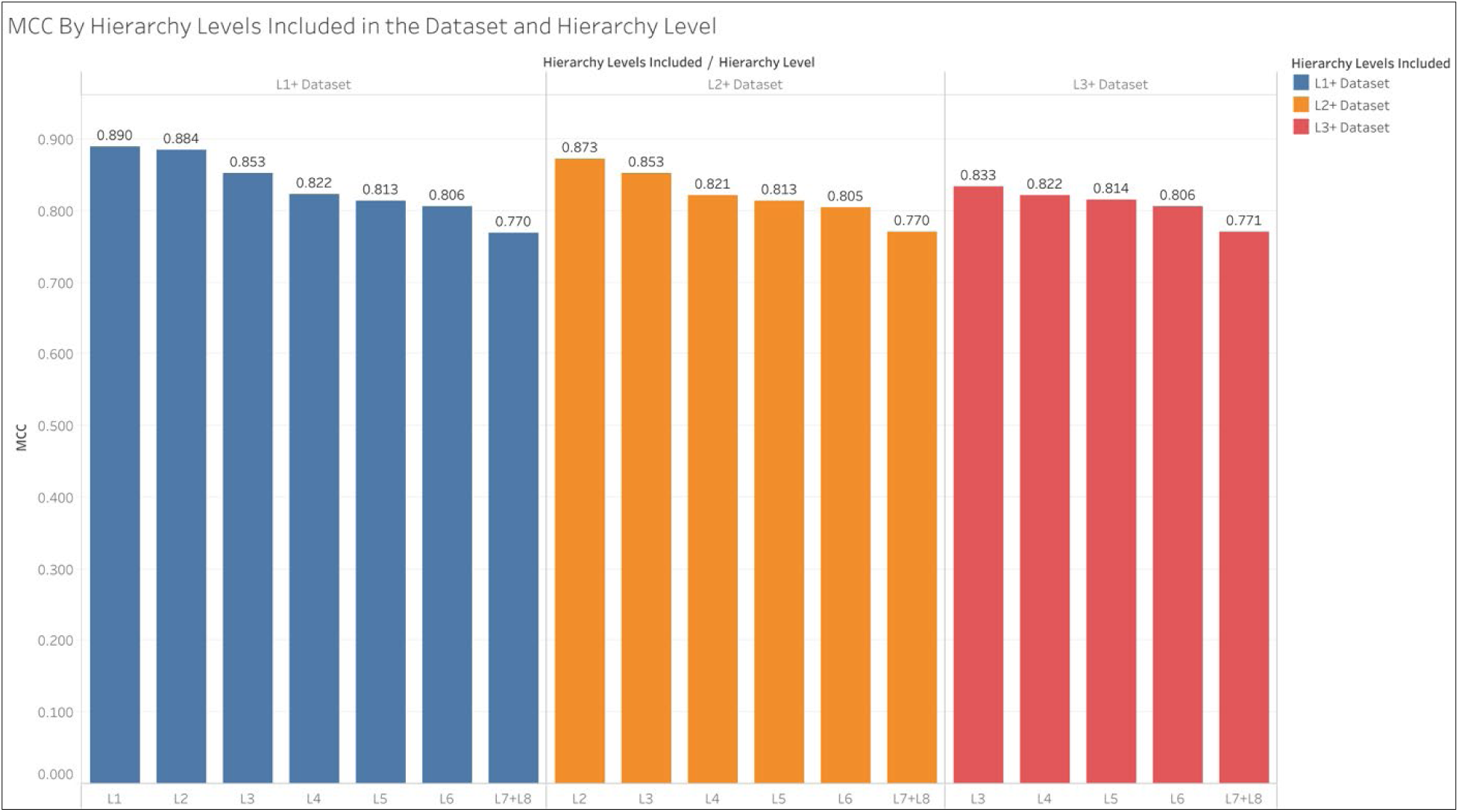
Comparing the overall MCC of pathway hierarchy levels within each dataset.

Fig 4 shows the same information as Fig 3, but compares performance across the datasets within hierarchy levels rather than the performance of hierarchy levels within the datasets. We see that inclusion of the L1 pathways in the L1+ dataset improved the performance of the L2 pathways. Similarly, inclusion of the L2 pathways in the L2+ dataset improved performance of the L3 pathways, though inclusion of the L1 pathways did not provide further improvement of the L3 pathways. Meanwhile the pathways in the L4, L5, L6, and L7+L8 hierarchy levels did not experience significant change in performance regardless of whether the L1 or L2 pathways were included in training.

**Fig 4.**
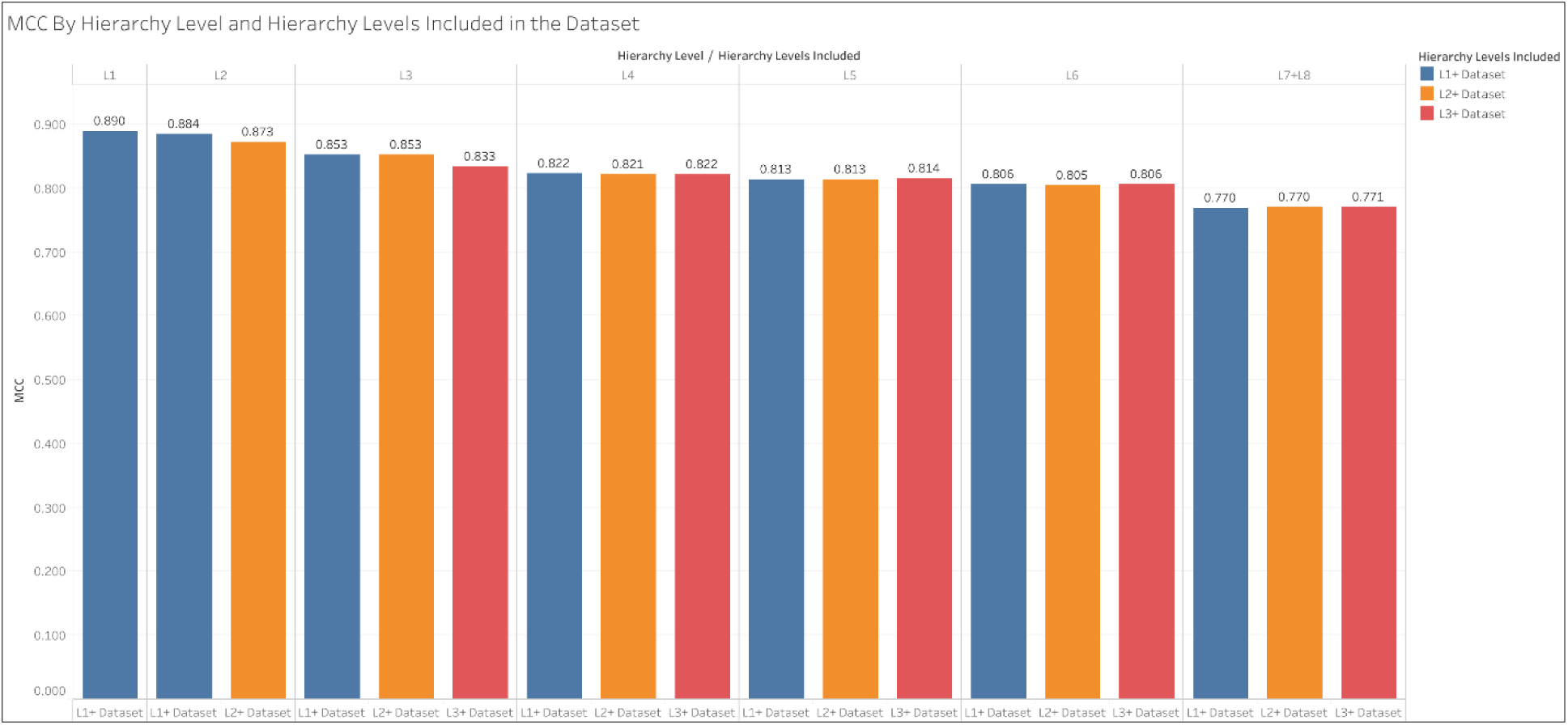
Comparing the overall MCC between datasets within each hierarchy level.

### MCC and compound/pathway size

We define compound size as the number of non-hydrogen atoms in the compound. We define pathway size as the sum of the sizes of the compounds associated with that pathway. Fig 5 shows the distribution of the size of compounds and pathways in the full MetaCyc dataset.

**Fig 5.**
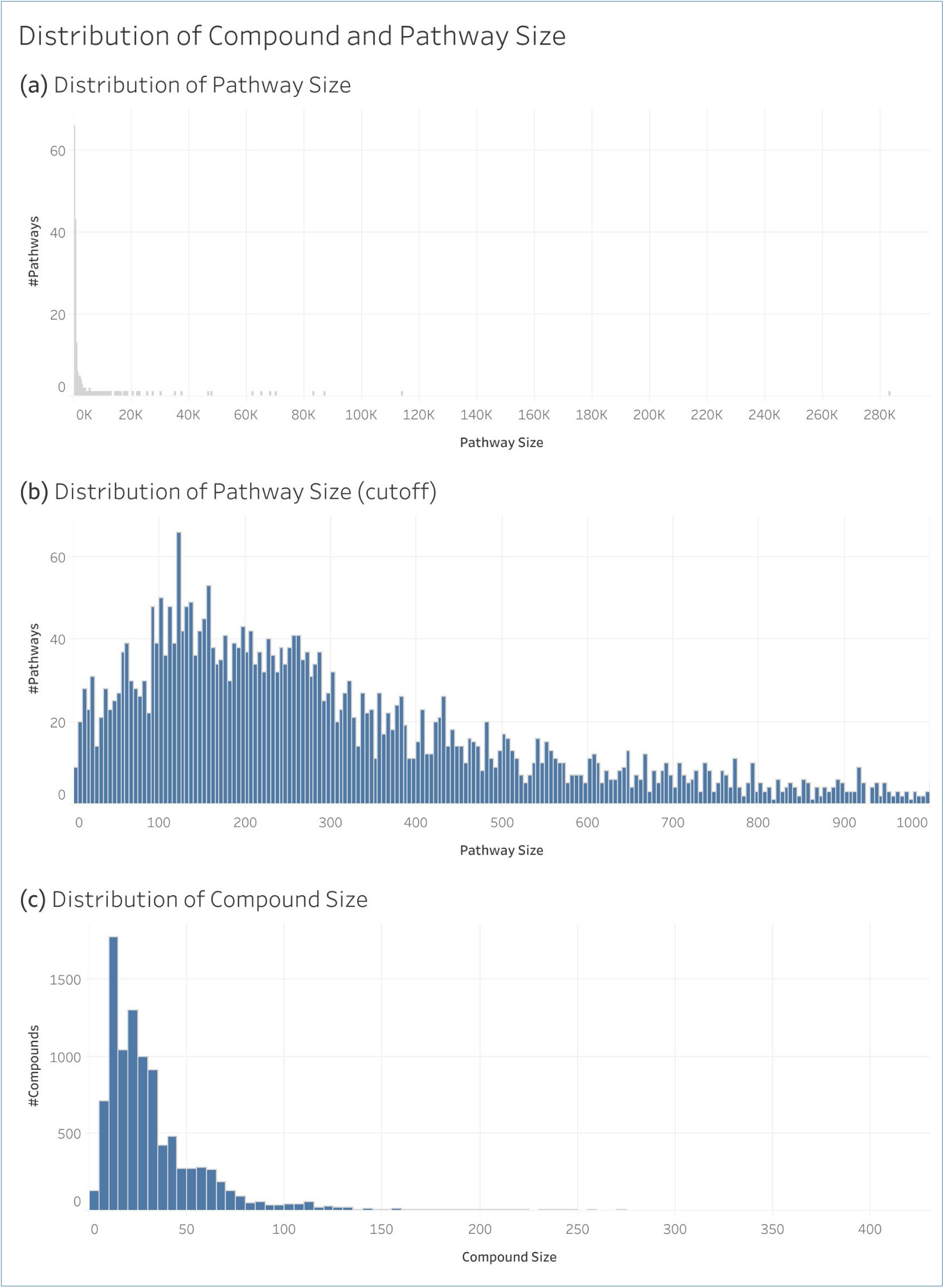
Distribution of compound size (number of non-hydrogen atoms in the molecule) and pathway size (sum of the sizes of compounds associated with a pathway).

### Distribution of Compound and Pathway Size

While Fig 5 shows the distribution of the size of all pathways, Fig 6 shows the distribution of pathway size within each hierarchy level. We see an overall trend of pathway size declining deeper into the hierarchy. Note that the y-axis is on a log scale to illustrate the wide range of pathway sizes that span multiple orders of magnitude.

**Fig 6.**
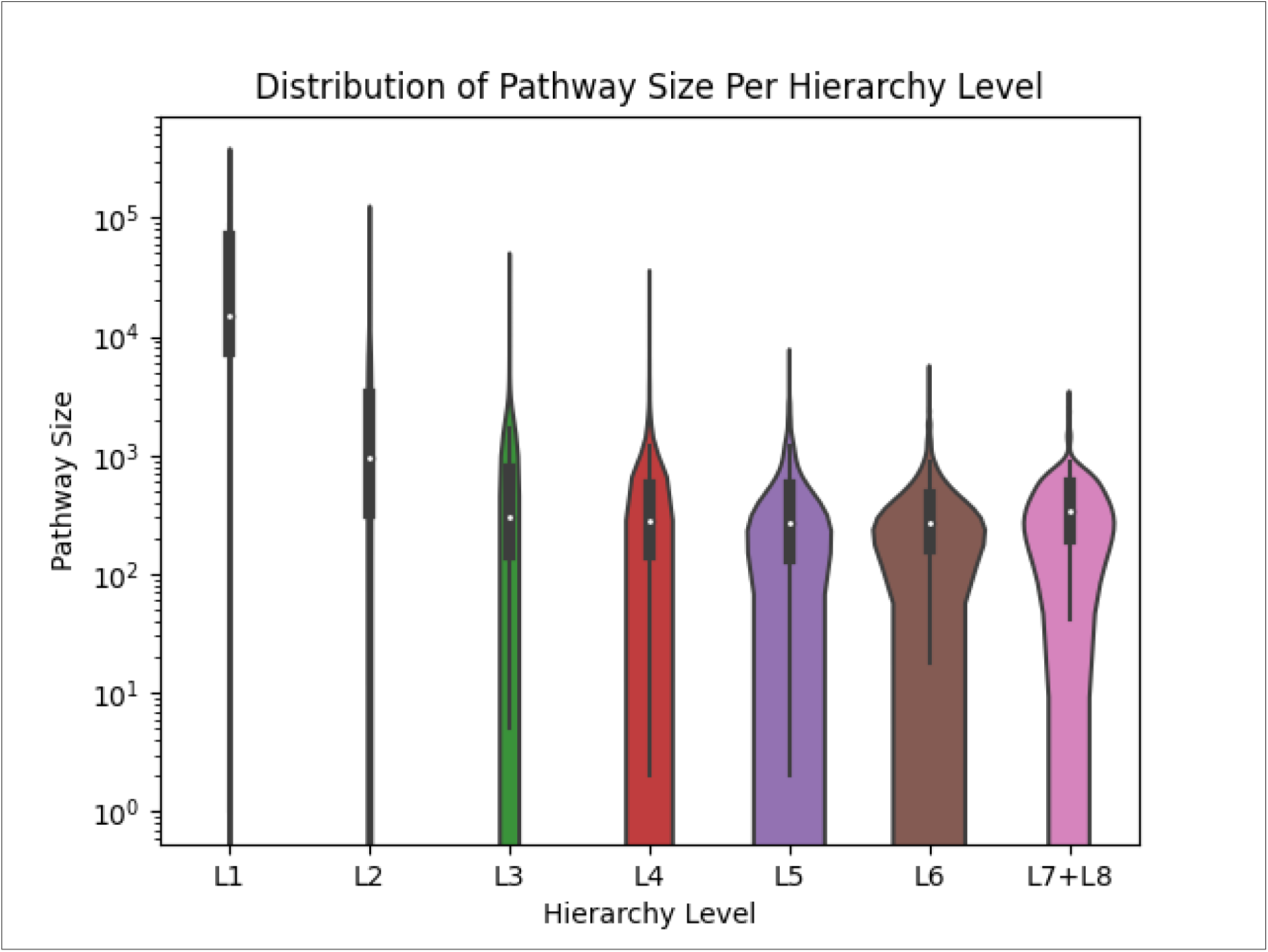
Violinplot showing the distribution of the sizes of the pathways within each hierarchy level.

Fig 7 shows the distribution of the overall MCC of individual pathways and compounds. There were 9,847 compounds and 4,055 pathway entries in the full MetaCyc dataset. The MCC distributions are negatively skewed for both pathways and compounds, with the compounds highly negatively skewed, likely due to truncation effects from boundary conditions.

**Fig 7.**
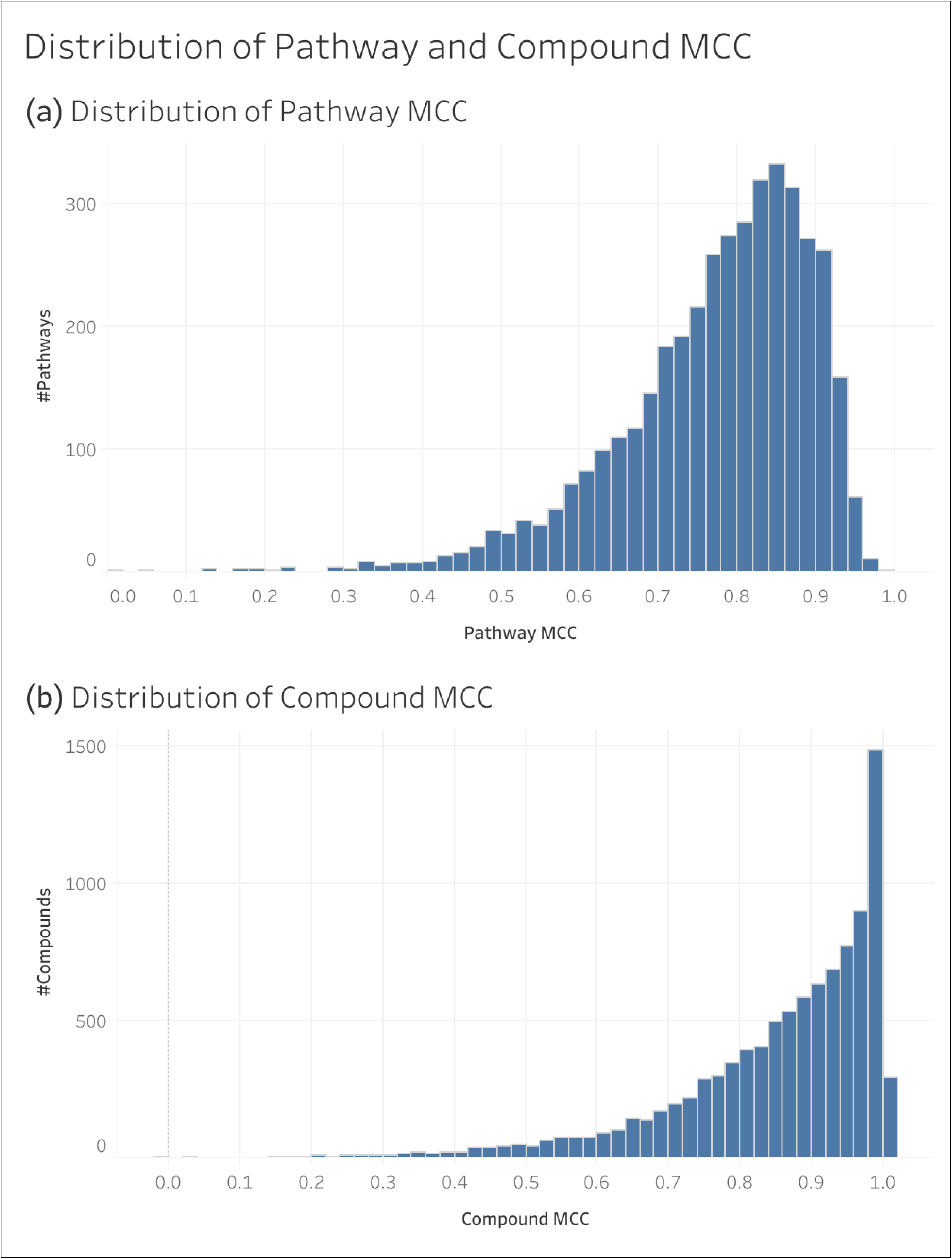
Distribution of the MCC of individual pathways and compounds.

#### Distribution of Pathway and Compound MCC

Fig 8 contains scatterplots comparing the MCC of pathways and compounds to their size. Fig 8a and Fig 8c compare pathway size to MCC and compound size to MCC respectively, while Fig 8b and Fig 8d are the same plots but with the x axis on a log scale for better visibility. We see that the pathway to MCC plot exhibits a funnel shape indicating that the variance of pathway MCC decreases as pathway size increases. Additionally, we see that the maximum MCC is lower for smaller pathways. Likewise, the maximum compound MCC does not reach 1.0 until reaching a compound size of 7 and we see before that, maximum compound MCC decreases as compound size decreases.

**Fig 8.**
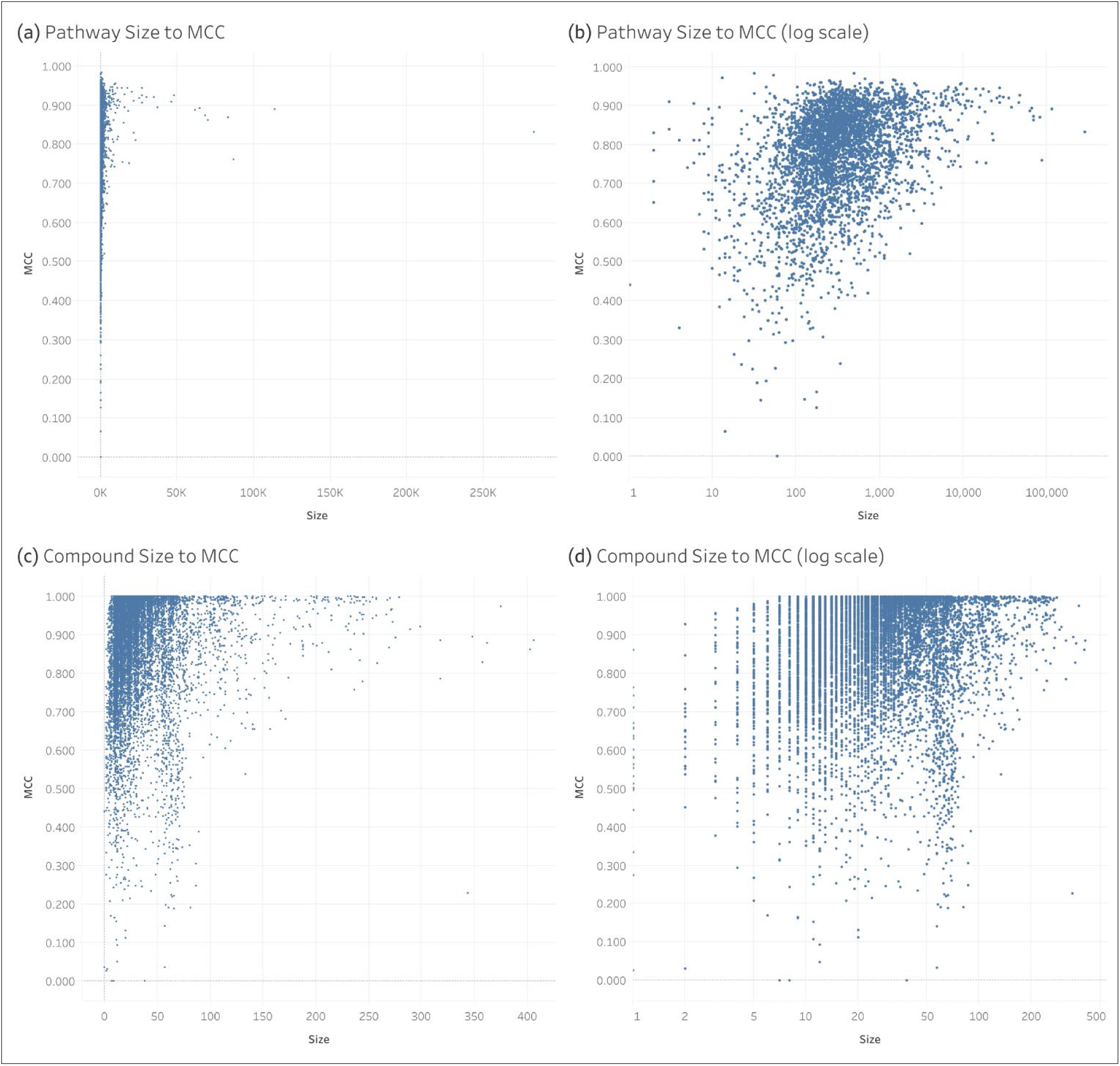
Scatterplots comparing the size of pathways and compounds to their MCC.

### Comparing MetaCyc to KEGG

Table 4 compares the performance of the full MetaCyc dataset, which primarily contains metabolic pathways, to the performance of the 184 KEGG pathways under the ‘Metabolism’ category as well as the full KEGG dataset containing all 502 pathways. We see that the KEGG metabolic pathways alone have the lowest mean MCC and the highest standard deviation. The full KEGG dataset has the highest mean MCC and lowest standard deviation. However, the difference in mean MCCs between the MetaCyc and full KEGG datasets is 0.002, representing a Cohen’s d of 0.2 which is considered to be a small effect size. Also, the percent mean difference is only 0.24%. Thus, the average performance of models trained on the MetaCyc dataset is comparable to that of models trained on the full KEGG dataset. However, if the comparison is limited to metabolic pathways in KEGG, then MetaCyc represents over a 5.6% improvement in prediction performance.

**Table 4.** Comparing the overall MCC of the MetaCyc dataset to past KEGG datasets.

| **Dataset** | **Mean MCC** | **Median MCC** | **Standard Deviation** | **Reference** |
| --- | --- | --- | --- | --- |
| **KEGG ('metabolism' pathways only)** | 0.800 | - | 0.021 | (8) |
| **MetaCyc** | 0.845 | 0.845 | 0.0101 | Current Study |
| **KEGG** | 0.847 | 0.848 | 0.0098 | (10) |

Fig 9 compares the distribution of MCC for the full KEGG and MetaCyc datasets. Visually, the variance and median are comparable for the KEGG and MetaCyc datasets, which is quantitatively corroborated in Table 4.

**Fig 9.**
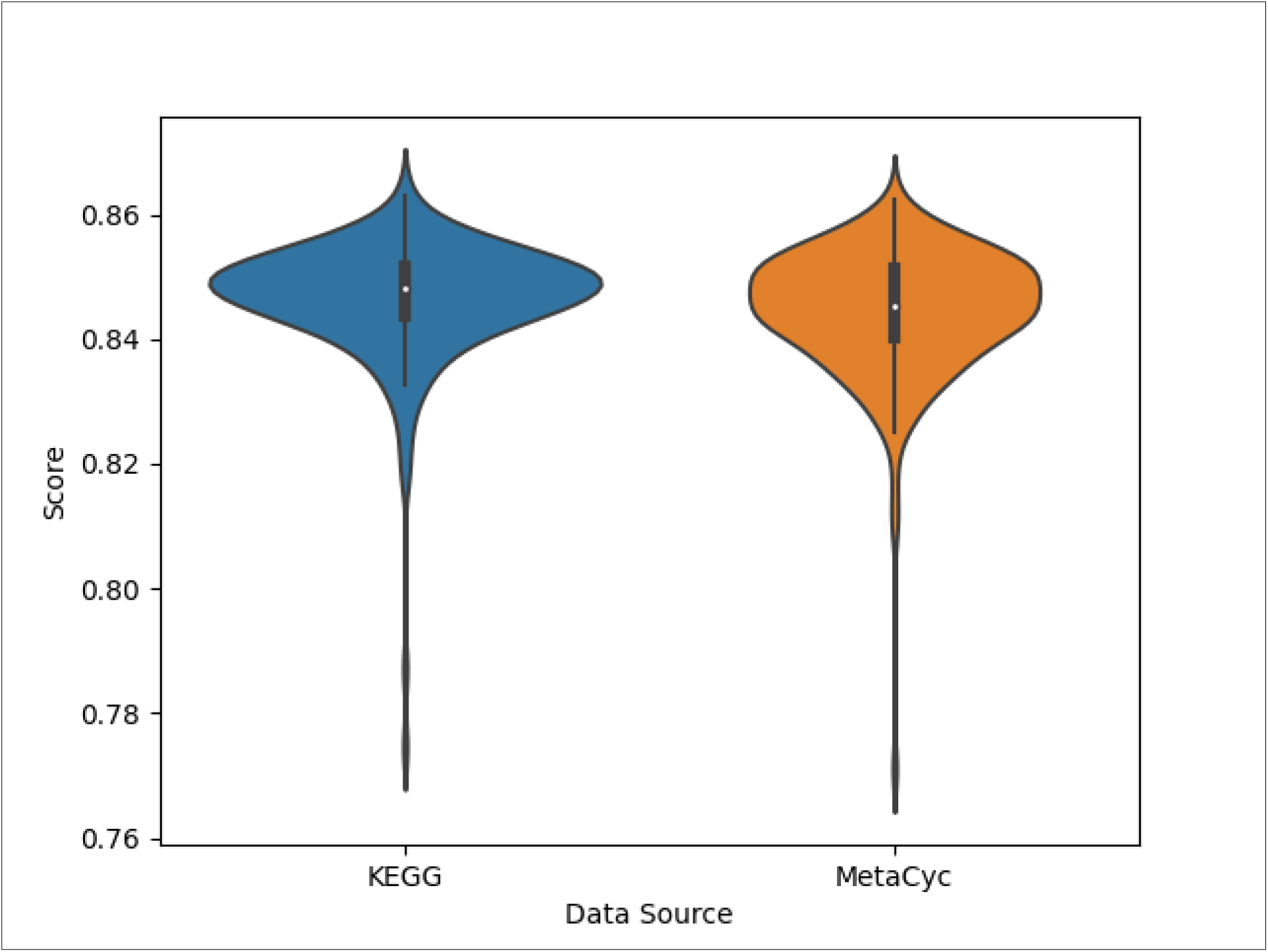
Violinplot comparing the distribution of the MetaCyc dataset’s MCC to that of KEGG.

Fig 10a shows the distribution of pathway size (sum of the number of non-hydrogen atoms across all associated compounds) in an overlapping histogram. Fig 10b shows the corresponding smoothed density plot. Metacyc clearly has a unimodal distribution, while KEGG has a bimodal distribution. KEGG has a higher proportion of larger pathways as indicated by the higher blue mode, while MetaCyc has several times the number of larger pathways, even though its proportion is smaller. MetaCyc also has a higher proportion of pathways with 100 to 1000 pathway size, indicated by the the single orange mode.

**Fig 10.**
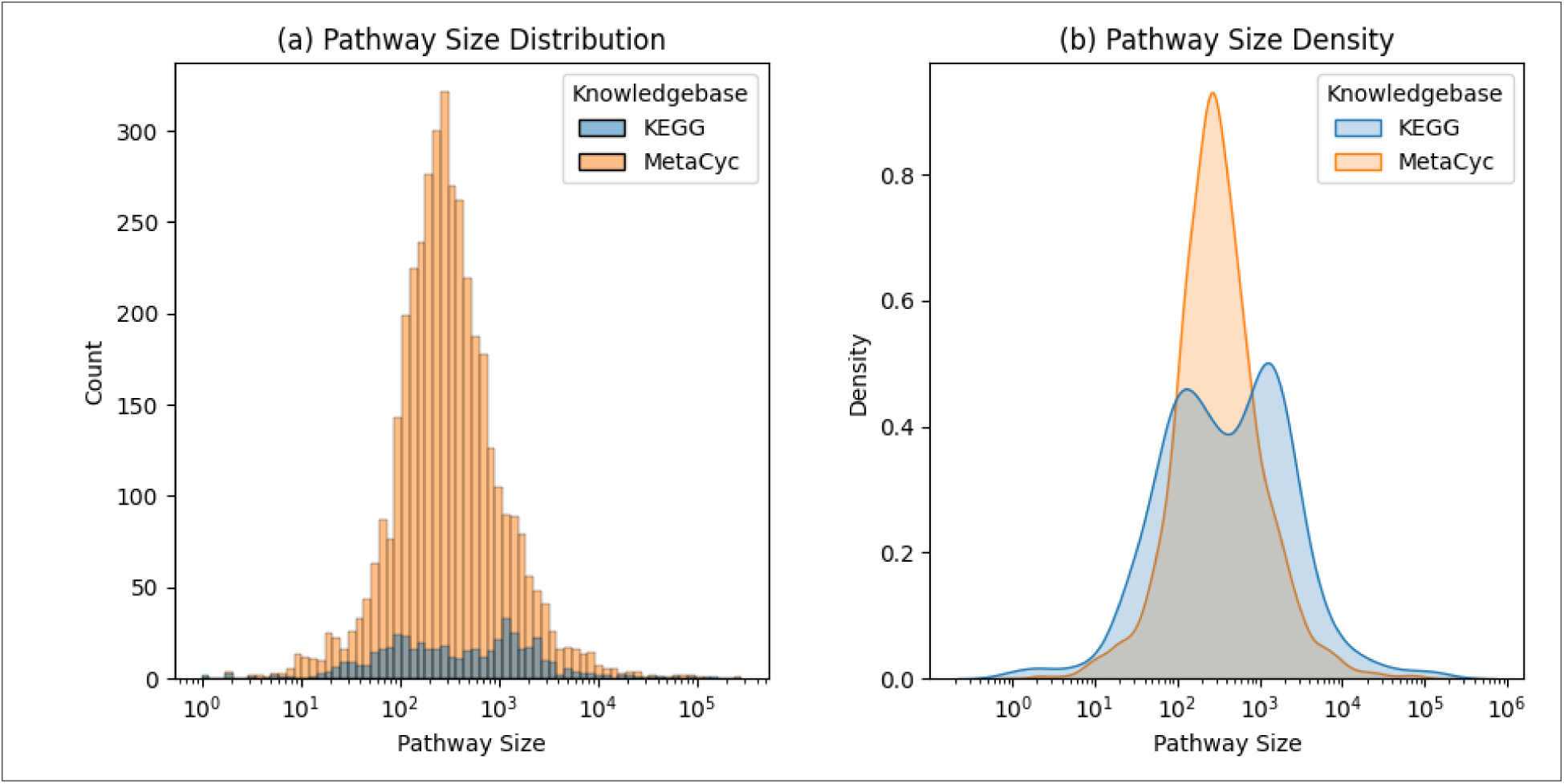
Distribution and density plot comparing pathway size between the MetaCyc and KEGG datasets.

## Discussion

While the majority of publications on pathway prediction have used KEGG data, the methods clearly are applicable to data from other knowledgebases. In this work, we demonstrate effective training with the MetaCyc knowledgebase for prediction of MetaCyc pathway involvement to a level comparable to the performance on the KEGG knowledgebase. There is less than a 0.24% difference in performance and the Cohen’s d effect size is only 0.2. Pragmatically, the overall performance is equivalent; however, if the comparison is limited to metabolic pathways in KEGG, then MetaCyc represents over a 5.6% improvement in prediction performance. Given the 10-fold larger size of the MetaCyc dataset, we may have reached asymptotic performance level of metabolic pathways with increasing dataset size. Moreover, the MetaCyc results expand the number of pathway definitions that can be predicted by 8-fold. The MetaCyc dataset also contains roughly 50% more compounds than the KEGG dataset, expanding the diversity of compounds available for model training and testing. Another key difference between the MetaCyc and KEGG datasets is the depth of the pathway hierarchy. KEGG pathways only span 3 levels, while MetaCyc pathways span 8 levels. The wider range of pathway granularity may have utility for biological and biomedical interpretation.

In addition, the results of the MetaCyc dataset are consistent with past results from the KEGG dataset pertaining to compound size, pathway size, hierarchy depth, and MCC. The deeper you go into the pathway hierarchy, the more granular the pathways become, having fewer associated compounds. The lower number of associated compounds results in smaller pathway size. Pathway performance declines with shrinking pathway size and with increasing depth in the pathway hierarchy. Additionally, the variance of pathway performance decreases as pathway size increases, suggesting that larger pathways have more robust prediction performance. We observe similar results with compound size, with the maximum MCC increasing as compound size increases. We recommend that researchers take compound and pathway size into account when using predicted pathway annotations.

The results of the MetaCyc dataset are also consistent with past results regarding the inclusion of higher-level pathways. Even if researchers are primarily interested in lower-level pathways, inclusion of higher-level pathways in the training set results in better performance of the lower-level pathways. Thus, we recommend that the full dataset be used to train a model, even if only a subset of the pathways is used in a downstream analysis.

## Acknowledgements

We thank the University of Kentucky Institute for Biomedical Informatics and National Science Foundation Grant Number 1626364 for their support and associated computing resources.

## Author Contributions

### Erik D. Huckvale

Obtained the data, wrote the code for the data engineering and analysis, finalized the results, and wrote the first draft of the manuscript.

### Hunter N. B. Moseley

Provided funding, worked in a supervisory role, reviewed the results, revised the manuscript, and provided mentorship.

## Data availability

All data and code for reproducing the results of this manuscript are available in the following figshare: https://doi.org/10.6084/m9.figshare.27317163.

## Supporting information

**Fig S1.**
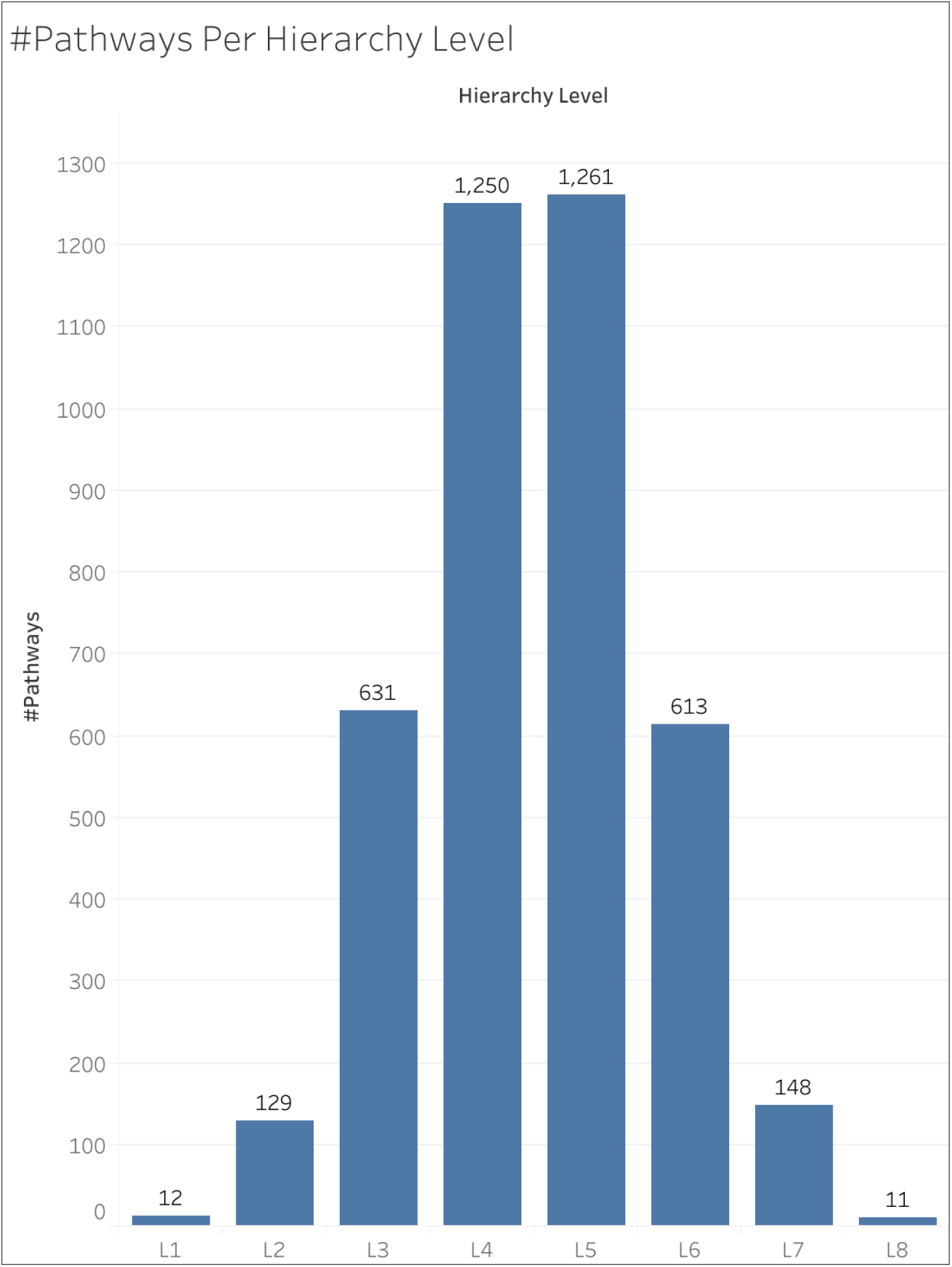
**The number of pathways within each hierarchy level.**

**Table S1.**
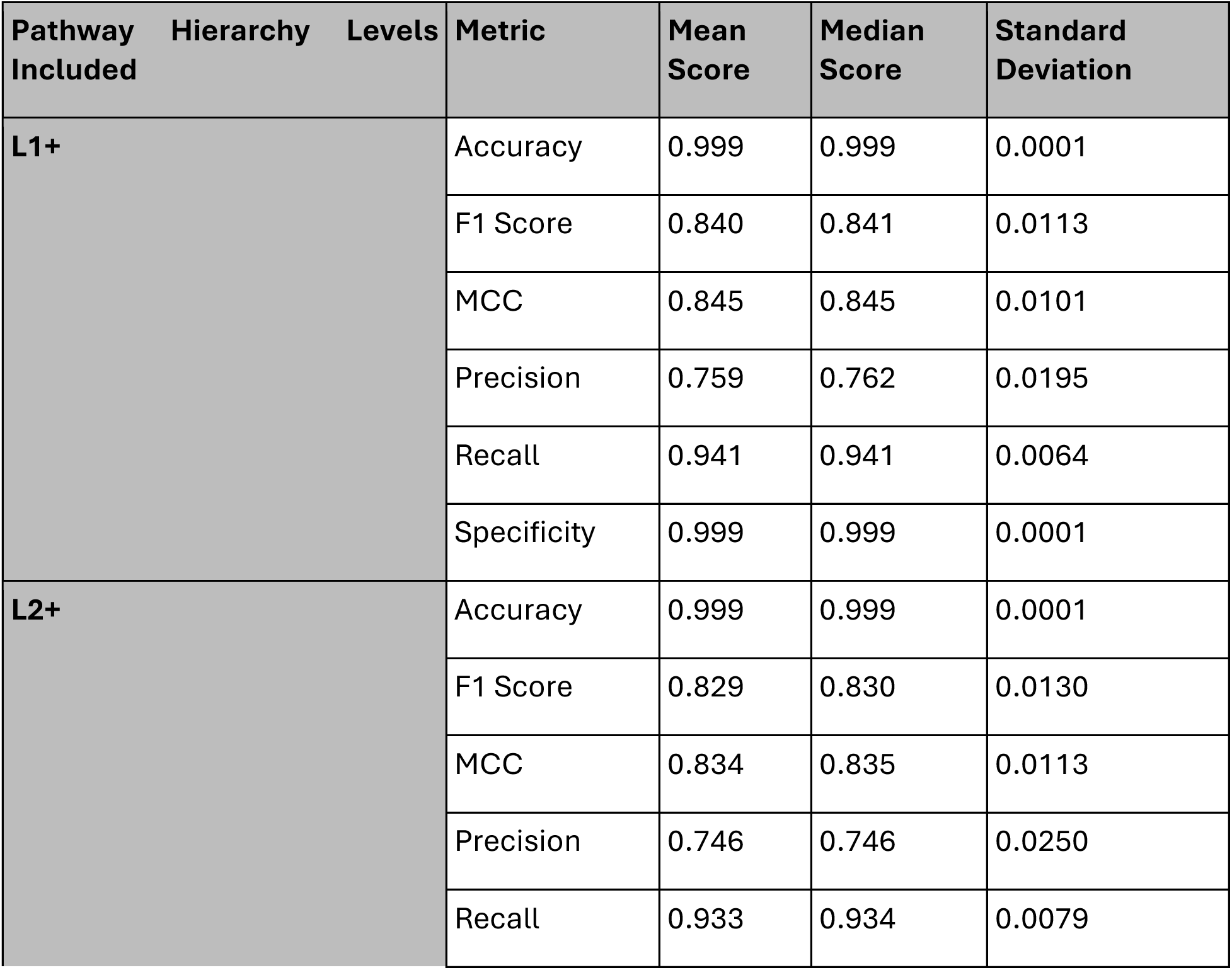
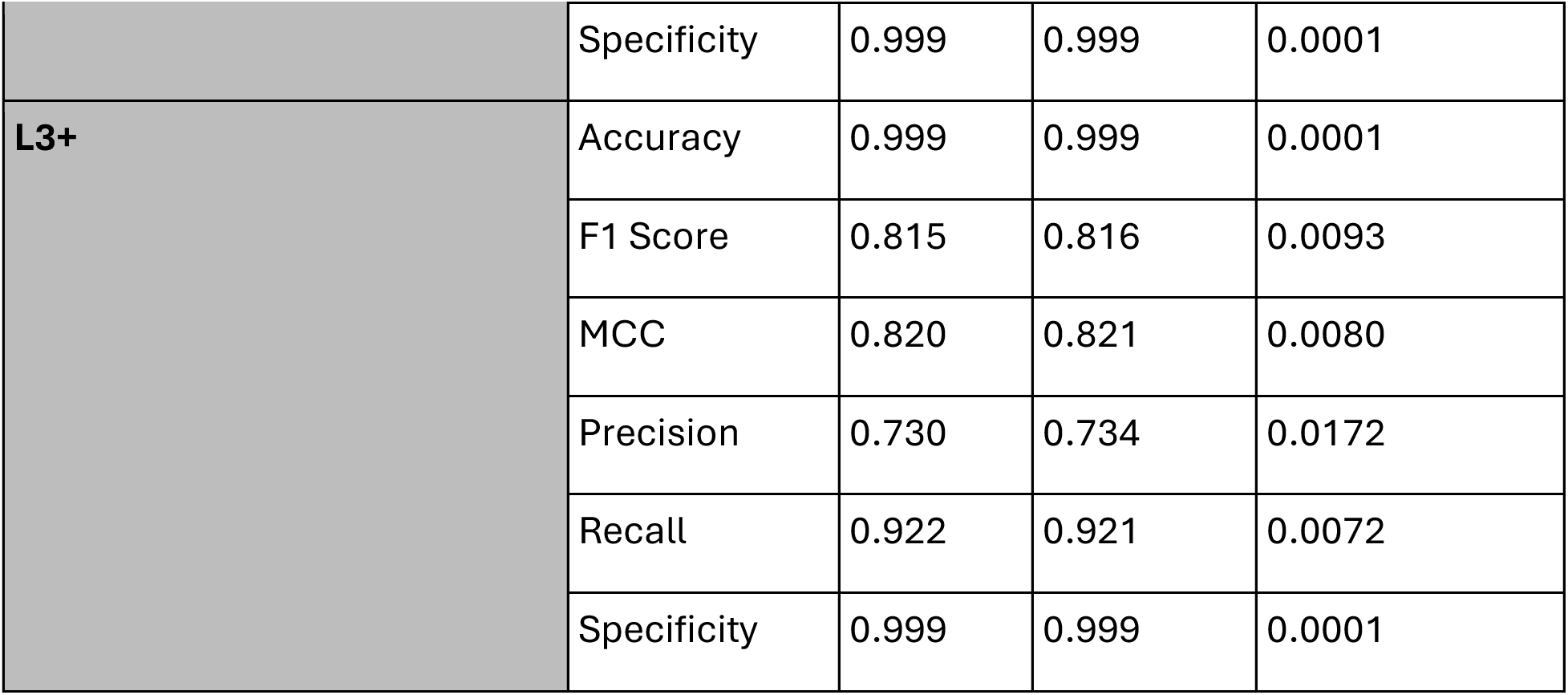
Statistics of performance metrics for models trained on the L1+, L2+, and L3+ datasets.

